# Differences in strategic abilities but not associative processes explain memory development

**DOI:** 10.1101/693895

**Authors:** N.C.J. Müller, N. Kohn, M. van Buuren, N. Klijn, H. Emmen, R.M.W.J. Berkers, M. Dresler, G. Janzen, G. Fernández

## Abstract

Children’s learning capabilities change while growing up. One framework that describes the cognitive and neural development of children’s growing learning abilities is the two-component model. It distinguishes processes that integrate separate features into a coherent memory representation (associative component) and executive abilities, such as elaboration, evaluation and monitoring, that support memory processing (strategic component). In an fMRI study using an object-location association paradigm, we investigated how the two components influence memory performance across development. We tested children (10-12 yrs., n=31), late adolescents (18 yrs., n=29) and adults (25+ yrs., n=30) of either sex. For studying the associative component, we also probed how the utilisation of prior knowledge (schemas) facilitates memory across age groups. Children had overall lower retrieval performance, while adolescents and adults did not differ from each other. All groups benefitted from schemas, but this effect did not differ between groups. Performance differences between groups were associated with deactivation of the dorsal medial prefrontal cortex (dmPFC), which in turn was linked to executive functioning. These patterns were stronger in adolescents and adults and seemed absent in children. This pattern of results suggests the children’s executive system, the strategic component, is not as mature and thus cannot facilitate memory performance in the same way as in adolescents/adults. In contrast, we did not find age-related differences in the associative component; with activity in the angular gyrus predicting memory performance systematically across groups. Overall our results suggest that differences of executive rather than associative abilities explain memory differences between children, adolescents and adults.

## 1. Introduction

In virtually all contexts learners need to focus on what to learn, avoid distraction, relate information to each other or keep several types of information online to combine them. These abilities, executive functions, have been shown to strongly influence the efficiency of ones mnemonic system (Simons and Spiers, 2003). The maturation of the executive system – especially the prefrontal cortex – during adolescence (Bunge and Crone, 2009; Crone and Dahl, 2012; Luna et al., 2015) makes it an excellent candidate to support the development of learning. The relation of executive functions with associative memory processes have been formalised and extended in a model explaining age-related differences in episodic memory: the two component model of development (Shing et al., 2008, 2010). It postulates one associative and one strategic component with differential maturational trajectories. The associative component “refers to mechanisms of binding together different features of the memory content into coherent representations” (Shing et al., 2010). The strategic component “refers to control processes that aid and regulate memory content at both encoding and retrieval” (Shing et al., 2010). Whereas the strategic component is centred around the prefrontal cortex, the associative component is centred around the medial temporal lobe. However, the developmental interaction of the two systems and their underlying neurobiology are still poorly understood.

When we learn new information this usually involves prior knowledge. Almost nothing we learn is fundamentally new in all aspects but mostly relates to something we already know. This entails that when we form new memory representations, the different features that get integrated via the associative component of the two-component model also include prior knowledge. That prior knowledge benefits learning was first formulated by Bartlett (Bartlett, 1932) and Piaget (Piaget, 1936) in the context of schemas: Our knowledge is organised in schemas which can be used to readily assimilate new information about the world or provide a foundation that can be modified when we acquire new insight/perspectives. The idea of schemas had a strong influence on education and educational psychology (Thorndyke and Hayes-Roth, 1979). Throughout our life we continuously acquire, modify, or enrich schemas. This difference in scope of schemas available to children versus adults might explain developmental memory differences (Brod et al., 2013). Whereas adolescents and adults have a sophisticated net of knowledge spanning a large range of topics, children are still in the process of acquiring most of that. Thus, for new information the children might have fewer opportunities to relate new information to their schemas. On the other hand, children might be superior in building new knowledge structures of previously unconnected information due to their generally increased neural plasticity (Ismail et al., 2016).

Executive processes and the utilisation of schemas have both been linked to the prefrontal cortex, yet vary in their precise localisation (Miller and Cohen, 2001; Ridderinkhof, 2004; Tse et al., 2011; van Kesteren et al., 2012). Generally, the prefrontal cortex shows a protracted maturation trajectory, reaching a matured state only in the mid-twenties (Gogtay et al., 2004; Shaw et al., 2008). This relatively late maturation has previously been linked to the development of cognitive control (Luna et al., 2015). Based on that we tested three age groups that differ strongly in prefrontal maturation: children between the age of 10-12 years, 18-year-old late adolescents and adults over twenty-five years old. To measure memory performance we used a game like object-location memory task (van Buuren et al., 2014). A strength of this paradigm is that it has no verbal requirements, which would have favoured the older groups as verbal memory itself is still developing in children (Vakil and Blachstein, 2007). The schema was trained during the first part of our study so that all groups have the same level of prior knowledge available to facilitate learning.

## 2. Materials and Methods

### 2.1. Participants

Ninety right-handed native Dutch-speaking volunteers participated in this study. As we investigated developmental differences related to differential maturation of the prefrontal cortex we tested three different age groups: Thirty adults aged between 25-32 years old (M = 26.9 years, SD = 21.9 months, 12 male), twenty-nine adolescents aged 18 (M = 18.5 years, SD = 3.1 months, 10 male) and thirty-one children aged between 10-12 years old (M = 11.0 years SD = 8.8 months, 8 male). All subjects had normal hearing and normal or corrected-to-normal vision. All participants were required to have no history of injury or disease known to affect the central nervous system function (including neuropsychological disorders such as dyslexia, autism and ADHD) and to not have MRI contraindications. Assessment of these were based on self-reports by the participants. Adults and adolescents were recruited from the student population of Radboud University, Nijmegen, and from the surrounding community. Children were recruited through presentations and flyers at local schools. The study was approved by the institutional Medical Research Ethics Committee (CMO Region, Arnhem-Nijmegen). Written informed consent was obtained prior to participation from all participants who were at least 18 years old; for the children participating both parents signed the informed consent.

Of these 90 participants, 4 children had to be excluded (2 did not want to complete the study, 1 moved excessively in the scanner, 1 due to an experimenter error); 2 adolescents were excluded as they did not complete the training at home; 1 adult had to be excluded due to an experimenter error. Of these 83 participants that completed the study, we excluded 11 (6 children, 1 adolescent, 4 adults) participants for the analysis based on their poor performance – see the fMRI data analysis section for details. All analysis focussed on this final set of 72 participants (21 children, 26 adolescents and 25 adults).

### 2.2. Summary of Procedure

The study spanned eight days in total. On day one participants came to the lab for a first session. The next four days they performed additional sessions at home. On day eight they returned to the institute for the final session. As paradigm we used an adapted version of the memory game task that was used in another study (van Buuren et al., 2014). Details of the paradigm are explained below. As additional measures we utilised a short verbal memory task, a fractal n-back task (Ragland et al., 2002), the Wisconsin Card Sorting task (WCST) (Heaton et al., 1993) and the forward digit span task (Alloway et al., 2008). The rationale for the additional measures is explained in a separate paragraph below.

On day one, participants came to the Donders Centre for Cognitive Neuroimaging and started in a behavioural lab with a practice of the n-back task, followed by the verbal memory task. Immediately after completing the verbal memory task, participants were taken into the MRI scanner where we first acquired a 10-minute resting state scan during which participants were instructed to lie still, think of nothing in particular and look at a black fixation cross on a white background. After that participants performed the n-back task and lastly, we acquired a structural scan. As the memory task required the use of a trackball– because of MRI-compatibility – participants had a practice session with the trackball (Kensington, Orbit Optical Trackball) that was used for all sessions of the memory task. During all uses of the trackball we instructed participants to operate the trackball with two hands: the right dominant hand moves the cursor and the left hand clicks.

After the practice, participants performed the first two sessions of the memory game. During the next four days, participants were instructed to “play” the memory game at home using a provided laptop and trackball. Participants were instructed to not skip a day and perform the task at roughly the same time of day. We monitored this online using the times the log files were created. Day six and seven were free of any experimental tasks. On day eight, the participants came back to the institute for the final session. The time of day during the two visits differed by maximally two hours to avoid time of day confounds. Day eight started with the final two parts of the memory task in the MRI scanner. Between the parts, which took both roughly 17 minutes, there was a break during which the participant could leave the scanner. As a last scan we acquired a second structural scan. Finally, we conducted the Wisconsin Card Sorting Test and the digit span in a behavioural lab. The total task time on day eight was around one hour.

### 2.3. Memory Paradigm

The task mimics the card game “memory/concentration” on two boards of cards and is adapted from (van Buuren et al., 2014) to make it more suitable for children. It is a 2×2 design (schema/no-schema x paired/new paired associates): one board was the schema while the other was the no-schema condition. Each board contained 80 objects in total. 40 of these objects were learnt during the first four days (paired associates). The remaining 40 objects (new paired associates) were added on day five and filled the remaining empty positions on each board (see Fig. 1 for an illustration of the task). On the schema board the place location associations stayed constant across the whole experiment, whereas on the other, the no-schema board, the associations were randomly exchanged every day. Due to this manipulation, participants learned the associations on the schema board over the course of four days; forming a schema that contains the object place association of the first 40 objects. Participants could utilise this schema when learning the second set – the new paired associations – of associations on day five for the schema board. The memory for all 160 associations across both boards was tested on day eight in the MRI scanner.

**Figure 1.**
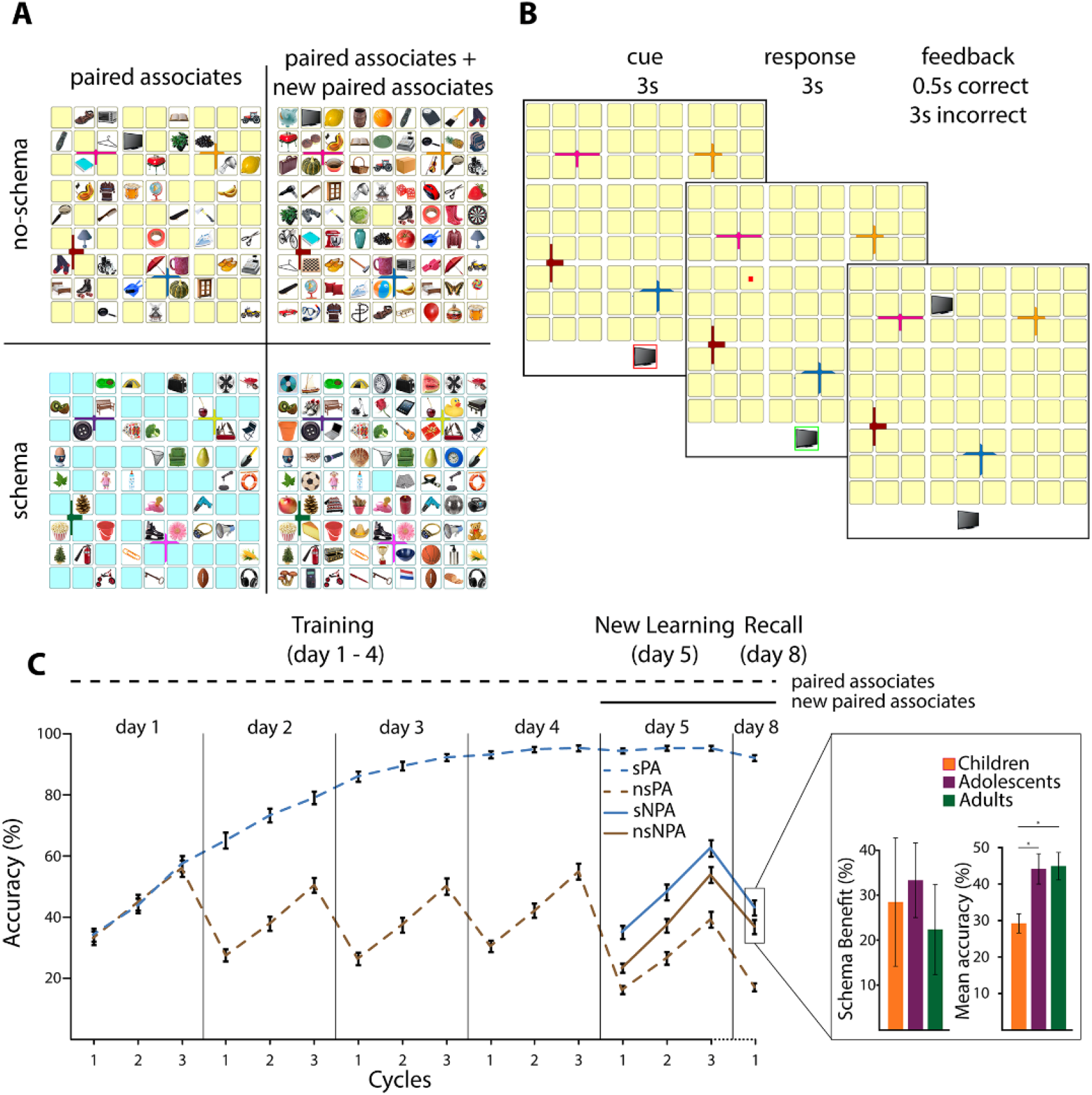
Task design and behavioural performance. **(A)** Participants needed to learn object-location associations (paired associates) in the memory game. For two boards (one schema, one no-schema board) there were two sets of associations to learn. During the first four days participants learned the paired associates on both boards (40 associations each). For the schema board participants could thus systematically learn the layout of the board. For the no-schema board on the start of every day the paired associates switched places with each other, therefore preventing systematic learning. On day five the new paired associates were added (again 40 associations per board). In the final session on day eight both the paired and the new paired associates were tested in a recall session in the MRI. **(B)** During a trial, participants first saw an object (cue) at the bottom of the board. After 3s the box in which the cue was presented turned green and a mouse cursor appeared. Participants then responded within 3s with the location corresponding to the object. If the response was correct the object was only shown very briefly (0.5s) whereas if they responded wrongly or not at all the object was shown for 3s. Each object was repeated three times for participants to have ample opportunity to learn the layout. Additionally, at the start of each session the whole board (during training the 40 paired associates, during new learning the whole 80 associations) Was presented. **(C)** During the training phase participants systematically learned the schema paired associates (sPA) on the schema board whereas the performance on the no-schema paired associates (nsPA) on the other board dropped at the start of every day due to the shuffling of locations. The schema new paired associates (sNPA) that were added during the new learning were better learned compared to the no-schema new paired associates (nsNPA) (F(1,69)=59.94, p<.001,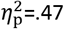). In the recall on day eight we observed a reduced performance in the children compared to both older groups (F(2,69)=5.33, p=.007, 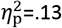). Schema benefit refers to how many items participants had correct in the schema new paired associates over the no-schema new paired associates. All error bars indicate the standard error of the mean. A star indicates a significance of p<.05.

### 2.4. Stimuli design, randomisation and presentation

In total we used pictures of 160 everyday objects. To ensure that especially the children could name all the objects effortlessly we selected the objects from a larger set of objects by asking an independent sample (n=5) of younger children (below the age of 9) to name all the objects. Only objects all children were able to name were included in the set of 160 pictures. The objects were randomly distributed across the different conditions (paired associates or new paired associates on the schema or no-schema board). This randomisation was done individually per participant.

Differing from the initial paradigm (van Buuren et al., 2014) we did not use a 10×10 board but a 9×9 board, furthermore we arranged the 9×9 board into nine visually segregated 3×3 boxes, each containing nine cards, by increasing the spacing after each third row and column. These changes had two reasons. First, we aimed to reduce the difficulty and the time required for the task to make it more suitable for children. Second, we opted to have an additional, more sensitive, measure of memory: instead of only taking into account the objects where the chosen position was exact, we can also analyse objects where the response was in the right box. Thus, we would be able to pick on memories where only an approximate location can be recalled. The two boards were differentiated by the colour of the back of the cards that were placed on the board. Whether the schema board or the no-schema board was yellow or blue was randomly assigned per participant in a counterbalanced fashion for each group separately.

We randomised the coordinates of the cards in a pseudorandom fashion separately per participant. Each 3×3 box of cards contained either four or five objects per condition (schema and no-schema) to ensure the cards were spread evenly across the board and there was no particular clustering. Furthermore, within each box there could not be a row of three objects, preventing particular easy structures from appearing within the box.

All of the memory tasks were implemented using Presentation version 18.1 (Berkeley, CA: Neurobehavioural Systems). The task at home utilised its web-based license. For tracking compliance with the study protocol, we used Dropbox (San Francisco, CA: Dropbox Inc.) on the laptops to automatically receive the log files. This communication was encrypted with a key only available to the researchers of this study to guarantee participants’ privacy.

### 2.5. Day 1: Basic training (in the lab)

Participants started by learning 40 paired associates on one board. The task started with a 1.5min presentation of all the 40 objects on their respective location to increase learning speed. After this initial phase the main task started and only the empty board was visible. One trial consisted of a cue-, a response- and a feedback phase: Participants saw an object at the bottom of the board in a red frame as cue. After 3s this frame turned green and a cursor appeared at a random position of the board. To draw attention to the location of the cursor there was a short animation (around 120ms) when it appeared. Participants now had to click within 3s on the location that belonged to the cued object using the trackball. When the response was correct the object was shown for 0.5s at its location. If there was no or a wrong response the cursor turned red and the object was shown at the correct location for 2.5s. After 40 trials there was a self-paced pause with a black fixation cross being presented instead of the board. The task consisted of three cycles. During each of those cycles every object was presented exactly one time. Participants therefore had three full training cycles for all items. After the three cycles were completed for the first board, the same procedure was repeated for the second board. Together both sessions took roughly 35min. We used the laptops and the trackball they would also use at home, showed them how to start the task and explained that the laptop needs an internet connection. To ensure understanding we had participants start the second part of the task themselves. The order of the boards and their colours were randomised and counterbalanced across participants per group.

### 2.6. Day 2-4: Training (at home)

During each of these three days participants would perform training sessions at home. For all sessions from now onwards the boards were presented in an interleaved fashion. During the initial encoding phase, first the one board and its 40 objects was presented for 1.5min, then the other board was presented for 1.5min. During the task, the board switched every five trials. This interleaved learning was used to reduce interference between the boards (McClelland et al., 1995). The start condition was randomised and counterbalanced across participants. Every day the associations on the no-schema board were shuffled as described above, preventing learning across days. Participants were instructed to try as hard as possible to perform well at that board and we showed performance scores at the end of the task to motivate them.

On day four, after the task, the training of the 80 paired associates was concluded. Immediately afterwards, participants performed a recall task testing all the paired associates they had learned. The trial structure for the recall was identical to the training except that there was no feedback and there was only one cycle: each object was shown once. The purpose of the recall task was two-fold; first to have a measure how well the paired associations were learned up until now; second, to familiarise the participants with the recall task before they would do the final recall in the scanner on day eight. Each session took around 35 minutes with day four being roughly five minutes longer.

### 2.7. Day 5: New learning (at home)

On day five the 80 new paired associates were added, 40 to each board. As before, the no-schema board was shuffled. The session started as usual with an initial encoding phase, however, now all 80 objects per board were shown and participants had 3min per board to memorise as many associations as possible. Aside from the number of associations, the session was identical to the previous training sessions. Each of the 80 objects per board was presented once per cycle leading to 480 trials in total. The boards were again presented in an interleaved fashion and the randomisation was done in such a way that there were never more than two paired associates trials or new paired associates trials in a row. The whole session took approximately 70 minutes. To reduce effects of exhaustion, participants were instructed to take a more prolonged self-paced break after 240 trials by standing up and moving around in the room, before resuming.

### 2.8. Day 8: Recall (in the MRI)

Around 72h later participants returned to the lab for the final recall in the MRI scanner. Participants lay down in the MRI scanner with the trackball positioned on their abdomen or their right upper thigh at a comfortable distance. Participants familiarised themselves with using the trackball in the scanner. After the participant was proficient using the trackball, we started with the recall task. One trial started as usual with a cue for 3s, followed by a response window of 3s followed by an inter-trial interval with only a black fixation cross on the screen for 2.5-7.5s. The inter-trial interval was drawn from a uniform distribution. There was no feedback presented during recall. To keep the trial length and the visual input consistent across subjects the board would still be presented for the whole duration of the response window, independent of whether the response was already given. The boards were again interleaved every five trials. We split the task in two parts (balanced across conditions) so that participants could take a break from scanning. Each part took roughly 17 minutes.

### 2.9. Additional Measures

Additional to the memory game we also conducted a short verbal memory task (day one, before scanning, outside the scanner), a fractal n-back task (Ragland et al., 2002) (day one, in the scanner), the WCST (Heaton et al., 1993) (day eight, after scanning, outside the scanner) and a forward digit span task (Alloway et al., 2008) (day eight, after scanning, outside the scanner). The verbal memory task was used to investigate links between cortical thickness of the prefrontal cortex and verbal memory performance. The fractal n-back was planned as a control experiment for a planned model-free analysis of the memory game. The WCST was included as an established measure of executive function. Finally, we included the digit span measure to control for group differences unspecific to long term memory processes.

### 2.10. Behavioural analysis

All statistical analyses of behavioural data were conducted in SPSS 21 (Armonk, NY: IBM Corp). Memory performance was measured as the number of correct responses per condition. The memory game consisted of four phases: training (day one till day four), recall (day four), integration of the paired associates (day five) and integration the complete set (day eight). The phases were analysed separately. To analyse the training (day one till day four) we used a repeated measures ANOVA with the factors schema (schema, no-schema), session (training day one to four), cycle (one to three) and group (children, adolescents, adults). For the recall on day four, the repeated measure model included schema and group. For the integration on day five we used a repeated measure model with the factors schema, cycle, group and included only the new paired associates. The model for the recall on day eight was identical to the recall on day four except that we now used the new paired associates instead of the paired associates. Whenever necessary, results were followed up with simple effect tests.

Complementarily, we repeated the analysis using the score for when participants clicked in the correct box (of the 9 boxes). This analysis is more sensitive, as responding close to the correct location likely also indicates memory; this heightened sensitivity comes at the cost of a higher chance level (11% versus 1.25%). We report only significant results with *p* < 0.05.

### 2.11. MRI data acquisition

Participants were scanned using a Siemens Magnetom Skyra 3 tesla MR scanner equipped with a 32-channel phased array head coil. The recall task comprised 935 volumes that were acquired using a T2*-weighted gradient-echo, multiecho echoplanar imaging sequence with the following parameters: TR = 2100ms; TE_1_ = 8.5ms, TE_2_ = 19.3ms, TE_3_ = 30ms, TE_4_ = 41ms; flip angle 90°; matrix size = 64 × 64; FOV = 224mm × 224mm × 119mm; voxel size = 3.5mm × 3.5mm × 3mm; slice thickness = 3mm; slice gap = 0.5mm; 34 slices acquired in ascending order. As this sequence did not provide whole brain coverage we oriented the FOV in a way that the hippocampus and the prefrontal cortex were fully inside and that only a small superior part of the parietal lobe was outside the FOV.

For the structural scans we used a T1-weighted magnetisation prepared, rapid acquisition, gradient echo sequence with the following parameters: TR = 2300ms; TE = 3.03ms; flip angle 8°; matrix size = 256 × 256; FOV= 192mm × 256mm × 256mm; slice thickness = 1mm; 192 sagittal slices.

### 2.12. MRI preprocessing

Preprocessing was done using a combination of FSL tools (Jenkinson et al., 2012), MATLAB (Natick, MA: The MathWorks) and ANTs (Avants et al., 2011a). From the two structural scans we generated an average using rigid body transformations from ANTs (Avants et al., 2011a), this procedure removed small movement induced noise. From the two scans and the average we always selected the scan with the least amount of ringing artefacts for all future analysis. If no difference was visible we used the average scan. These scans were denoised using N4 (Tustison et al., 2010) and generated a study-specific template with an iterative procedure of diffeomorphic registrations (Avants and Gee, 2004). For the registration of the functional volumes we resampled the created template to a resolution of 3.5mm isotropic. Using Atropos (Avants et al., 2011b) the anatomical scans were segmented into 6 tissue classes: cerebrospinal fluid, white matter, cortical grey matter, subcortical grey matter, cerebellum and brainstem. The segmentation also produced individual brain masks.

For the functional multiecho data we combined echoes using in-house build MATLAB scripts. It used the 30 baseline volumes acquired during the resting period directly before each part of the task to determine the optimal weighting of echo-times for each voxel (after applying a smoothing kernel of 3mm full-width at half-maximum to the baseline volumes), by calculating the contrast-to-noise ratio for each echo per scan. This script also directly realigned the volumes using rigid body transformation. Afterwards the volumes were smoothed using a 5mm full-width at half-maximum Gaussian kernel and grand mean intensity normalisation was done by multiplying the time series with a single factor. Younger participants tend to move more than older ones. For a developmental study it is thus important to minimise the effect of motion in the data. For this purpose we applied AROMA, a state of the art motion denoising algorithm that uses independent component analysis decomposition of the data to identify movement and other noise signals (Pruim et al., 2015b, 2015a). Variance in the BOLD signal that could only be explained by components identified in this manner was regressed out. Afterwards we regressed out signal stemming from the cerebrospinal compartments and from the white matter by extracting the signal from individual generated segmentations using ANTs (Avants et al., 2011b). As a last step a 100s highpass filter was applied.

Boundary based registration was first calculated from native functional to native structural space using FLIRT (Greve and Fischl, 2009). We then calculated nonlinear registration from native structural space to the study template with FNIRT (Smith et al., 2004). The warping was done in a way that every functional volume was only resliced exactly once after the initial realignment. For displaying the final results we warped the final maps to MNI space using the nonlinear registration of ANTs (Avants and Gee, 2004).

### 2.13. fMRI data analysis

After preprocessing the data was analysed using the general linear model framework implemented in FEAT (Jenkinson et al., 2012). On the first level we included eight separate regressors: four regressors modelled correct responses for the separate conditions (paired associates vs. new paired associates on the schema or no-schema board). As duration we used the trial onset of the cue until the participant gave a response. As a correct response, we counted if the participant clicked in the correct one of the nine boxes. We used this way of scoring instead of using only the trials in which participants clicked on the correct card as we would have substantially more power due to the higher amount of trials for the MRI analysis while still maintaining a fairly low chance level (11%).

For all those conditions but the schema paired associates an additional regressor was included to model incorrect responses. For the schema paired associates condition the performance was designed to be as close to ceiling as possible leading to only few incorrect trials. These trials were modelled together with all the trials in which subjects failed to respond in time in a single “miss” regressor. For the miss trials the full 6s of the cue and response window was used. Regressors were then convolved with a double gamma hemodynamic response function. On the first level the model was fitted separately per run. Using fixed effect modelling the runs were combined per subject and then the participant specific contrasts were estimated. To calculate the group level statistics, we warped the participant level results into study template space and used mixed effect modelling implemented in FSL FLAME2. The results were thresholded using a cluster forming threshold of *z* > 2.3 (equal to *p* < 0.01) and a cluster significance threshold of *p* < 0.05 at the whole brain level. Our central motivation for this study was to understand how the neural mnemonic processes differ across different stages of cortical maturation. Therefore, our imaging analysis was centrally guided by the behavioural results, illuminating the underlying neural architecture related to behavioural differences. Thus, the contrasts used will be explained while presenting the imaging results.

As a follow up analysis, we conducted moderation analysis based on the results of the general linear model analysis. For this, we extracted the average betas from the significant clusters on a participant by participant basis. We then conducted the moderation analysis using the PROCESS macro (Hayes, 2013) for SPSS.

## 3. Results

### 3.1. Training

As expected, during the recall on day four schema items were better recalled than no-schema items (F(1,69)=199.05, *p*<.001, 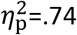). For the training, we observed a significant three-way interaction of schema x session x cycle (F(4.49,309.86)=27.84, *p*<.001, 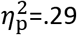), reflecting the fact that in the schema condition the paired associates could be learned across days while the shuffling between days prevented this for the no-schema paired associates.

### 3.2. New Learning

The schema new paired associates were learned better compared to the no-schema paired associates (F(1,69)=59.94, *p*<.001, 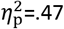).

### 3.3. Recall day 8

In the final recall the schema new paired associates were better retrieved compared to the new no-schema paired associates (F(1,69)=17.09, *p*<0.001, 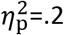). However, there was a significant effect of group on the retrieval performance of the new paired associates overall (F(2,69)=5.33, p=.007, 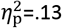). Children performed worse than adolescents (MD=−5.97, p=.006) and adults (MD=−5.23, p=.005); whereas adolescents and adults did not differ significantly (MD=−.31, p=.881).

### 3.4. Precise location correct vs. box correct

Analysing the data counting only trials as correct where the response was on the right card instead of the right box (3×3 cards) did not alter the results: All of the reported effects were also significant for the box score.

### 3.4. fMRI: Developmental differences

Our central behavioural finding is that children show lower memory performance than adolescents and adults while the latter two groups did not perform significantly differently. To understand the neural changes across development, we contrasted the activation during retrieval of the new paired associates for the correctly retrieved trials minus the trials in which a wrong response was given. As all groups seem to have profited to a similar degree from schema, we averaged across schema and no-schema trials to be more sensitive for developmental differences. The contrast between hits and misses was then compared between the children versus the average of the two older groups; this was done because the latter two groups did not differ in performance. As we used mixed modelling on the group level analysis the fact that the adolescent-adult group is twice as big as the children does not bias the results.

We observed increased activation in children in midline structures, including the dorsomedial prefrontal cortex (dmPFC). Increased activation in the adolescent-adult groups was most pronounced bilaterally in the angular gyrus (Fig. 2). For a complete list of all clusters please see Table 1.

**Figure 2.**
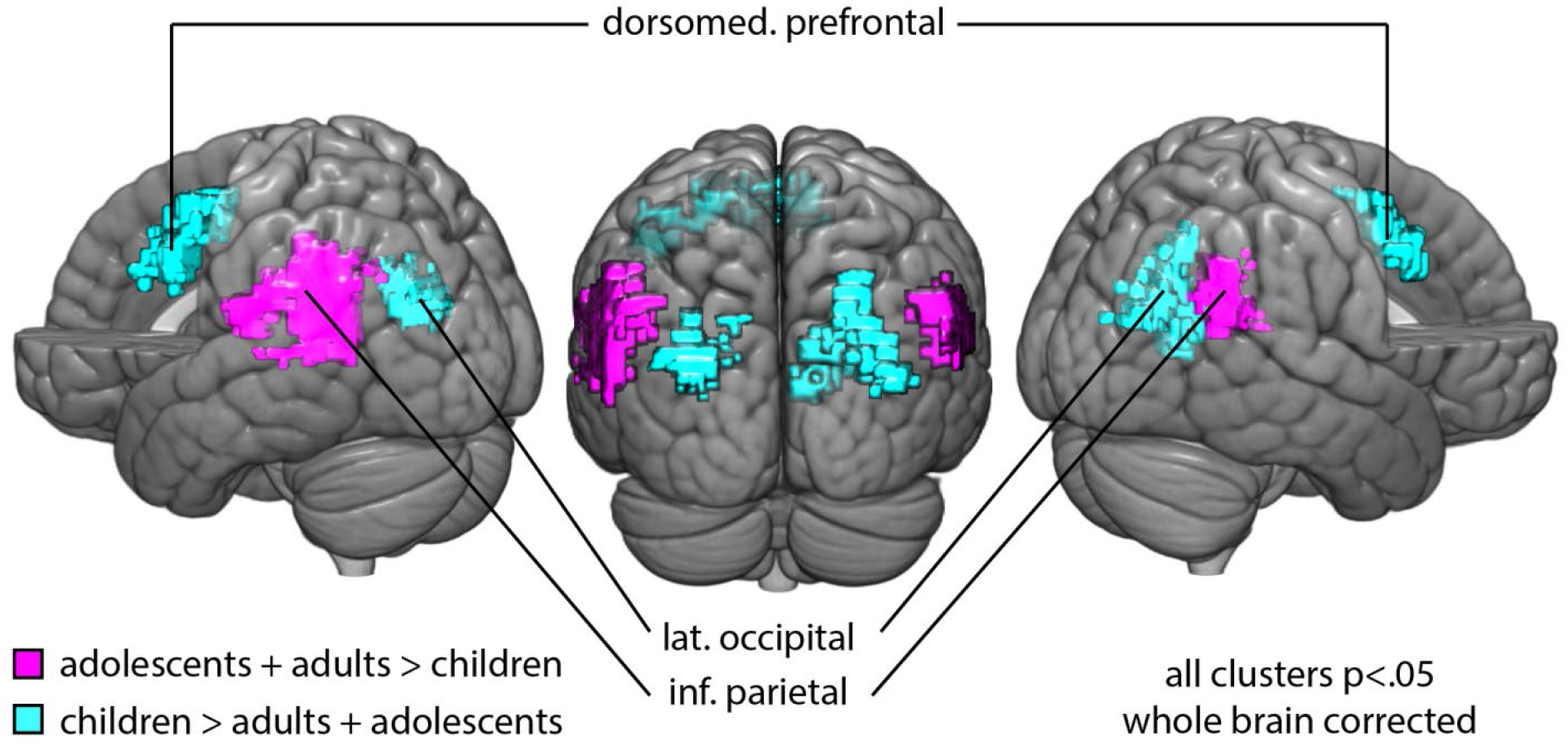
Age-related differences in mean memory performance for the new paired associates. During the recall of both the schema and the no-schema new paired associates children showed an increased activation in the dorsal medial prefrontal cortex (dmPFC) overlapping with the cingulate and paracingulate gyrus; a second cluster around the lateral occipital cortex showed the same effect. Adolescents and adults showed higher bilateral activation of the angular gyrus.

**Table 1.** Developmental differences in activation for the correct retrieval of the new paired associates. The listed clusters here and their local maxima show differences between the children, the adolescent and the adult groups for the retrieval of the new paired associates in which both older groups outperformed the children. The coordinates were always of the global/local maximum. The voxel-count as well as the z-score of the peak voxel were taken from study space. The MNI coordinates were obtained by warping the results into MNI space. All labels refer to regions on the cortex.

| Region | MNI Coordinates |  |  | z-Score | Voxels |
| --- | --- | --- | --- | --- | --- |
|  | x | y | z |  |  |
| Children > Adolescents + Adults |  |  |  |  |  |
| R dorsomed. prefrontal cort. | 3 | 15 | 46 | 4.04 | 243 |
| L dorsomed. prefrontal cort | -7 | -14 | 43 | 3.74 |  |
| R dorsomed. prefrontal cort | 4 | 22 | 42 | 3.72 |  |
| L precentral gyrus | -40 | 3 | 40 | 3.54 |  |
| R middle frontal gyrus | -25 | 2 | 55 | 3.48 |  |
| L dorsomed. prefrontal cort | -4 | 5 | 55 | 3.34 |  |
| R lat. sup. occipital cort | 25 | -67 | 41 | 4.73 | 151 |
| R precuneus | 15 | -58 | 25 | 3.5 |  |
| R lat. sup. occipital cort | 42 | -83 | 24 | 3.34 |  |
| R lat. sup. occipital cort | 30 | -80 | 29 | 3.3 |  |
| R Precuneus | 9 | -59 | 9 | 3.26 |  |
| R med. occipital cort | 38 | -76 | 19 | 3.17 |  |
| L lat. sup. occipital cort | -29 | -84 | 32 | 4.2 | 103 |
| L lat. sup. occipital cort | -26 | -81 | 24 | 3.9 |  |
| L lat. sup. occipital cort | -30 | -74 | 22 | 3.82 |  |
| L precuneus | -15 | -72 | 38 | 3.39 |  |
| L precuneus | -19 | -81 | 35 | 2.68 |  |
| L lat. sup. occipital cort | -45 | -85 | 27 | 2.54 |  |
| Adolescents + Adults > children |  |  |  |  |  |
| L lat. sup. Occipital cort | -54 | -61 | 44 | 4.89 | 227 |
| L angular gyrus | -51 | -59 | 32 | 4.14 |  |
| L lat. sup. occipital cort | -46 | -72 | 38 | 4.01 |  |
| L par. operculum | -63 | -34 | 22 | 3.85 |  |
| L lat. sup. Occipital cort | -58 | -64 | 16 | 3.61 |  |
| L mid. temp. gyrus | -69 | -48 | 6 | 2.83 |  |
| R lat. sup. occipital cort | 60 | -61 | 33 | 3.44 | 227 |
| R lat. sup. occipital cort | 48 | -61 | 42 | 3.34 |  |
| R lat. sup. occipital cort | 56 | -60 | 42 | 3.27 |  |
| R angular gyrus | 64 | -53 | 25 | 3.07 |  |
| R angular gyrus | 54 | -47 | 23 | 2.82 |  |
| R lat. sup. occipital cort | 60 | -61 | 21 | 2.8 | 103 |

As we hypothesised that performance differences might be due to differences in executive abilities in children, we tested whether there is a link between the (de)activation of the dmPFC and measures of executive function. Activation in the dmPFC was negatively correlated with performance in the WCST (r(70)=−.31, p=.008), as measured by the amount of correct categories, but not significantly related to the forward digit span score (r(70)=.04, p=.72). The correlation of dmPFC activation and WCST performance was driven by a negative correlation across the two adult groups (r(49)=−.31, p=.026). This association between dmPFC activity and WCST performance for the children was reduced compared to the adolescent-adult group (z=2.08, p=.038) and in itself not significant (r(19)=.26, p=.27).

### 3.5. fMRI: Schema effect

Participants across all age groups remembered schema new associates better than the no-schema new associates. To illuminate the neural architecture behind this schema effect we calculated the contrast between the hits and the misses between the schema new paired associates (sNPA) and the no-schema new paired associates (nsNPA): sNPA (hits – misses) – nsNPA (hits – misses). However, there was no significant activation that survived whole brain correction. More specifically, we tested the angular gyrus, as the region had previously been found to be important for integrating different parts of a schema (Wagner et al., 2015) and in the paradigm utilised here in same the schema x memory contrast (van Buuren et al., 2014). When separately contrasting sNPA (hits – misses) and nsNPA (hits – misses), we observed that both angular gyri were significantly activated in both contrasts (p<.05). To test whether there was activation specificity for schema, we extracted the betas for voxels that were significantly activated for the sNPA’s; we extracted both the values for sNPA (hits – misses) and nsNPA (hits – misses). The difference between those contrasts was positively correlated (r(70)=.34, p=.003) to the magnitude of the schema benefit. There was no indication that this relation is significantly modulated by age group (F(2,66)=.95, p=.39).

## 4. Discussion

We tested how differences in the associative and the strategic component of the two-component model of memory development (Shing et al., 2008, 2010) contribute to memory performance differences between children, adolescents and adults. We found that both the adolescent and adult group had higher memory performance than children, independent of the conditions, while all groups profited equally from utilising schemas (Fig. 1). Performance differences between groups were associated with deactivation of the dmPFC, which in turn was linked to executive function. In contrast, activation of the angular gyrus was consistently correlated with memory performance across all groups (Fig. 3). This suggests that age-related differences in memory are rather driven by differences in the strategic component, but not the associative component.

**Figure 3.**
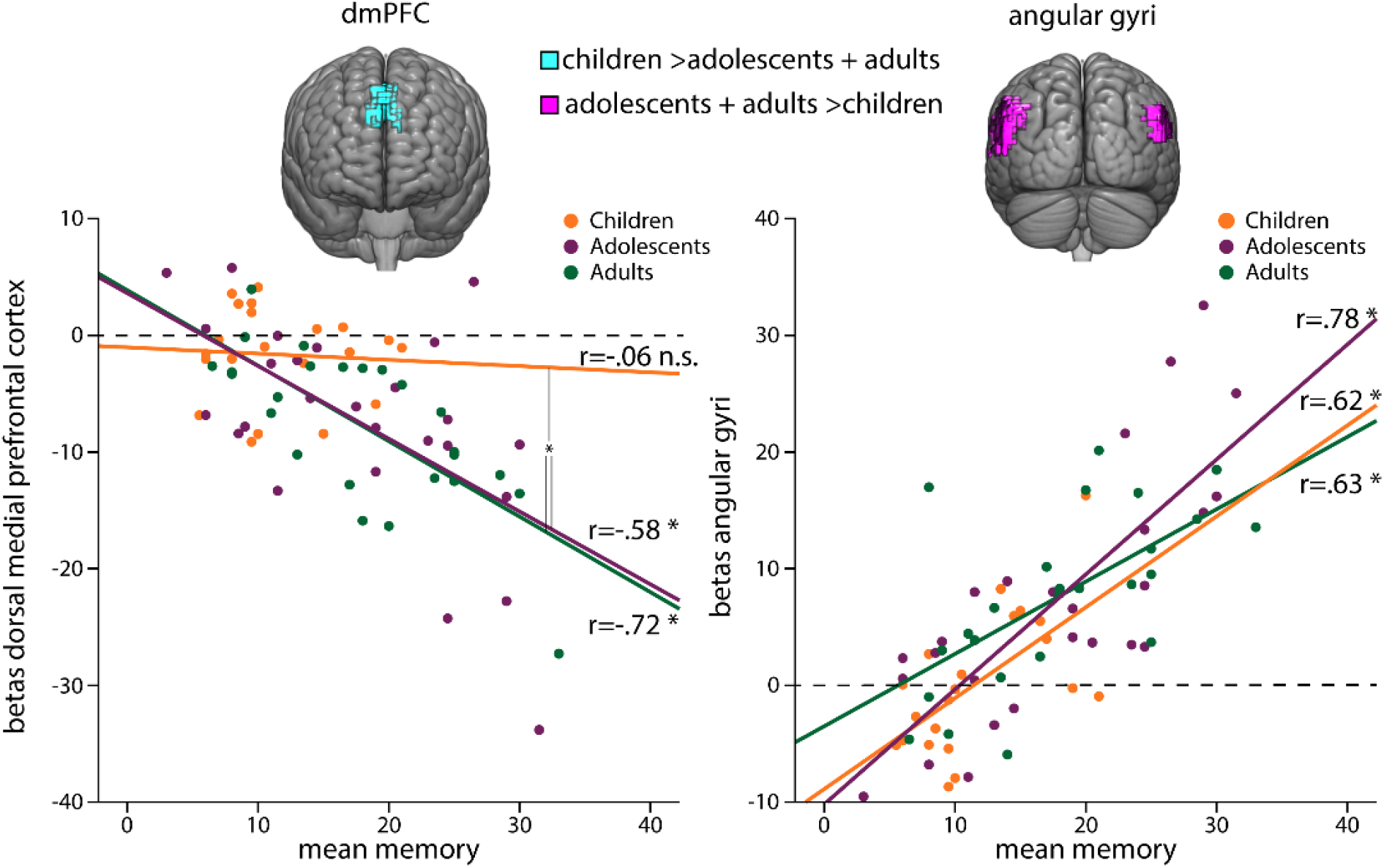
Developmental differences in brain-behaviour relation. For both the activation in the dmPFC and the angular gyri we found a relation to the mean memory performance across the age groups. For the dmPFC this relation was negative (r(70)=−.63, p<.001). For the angular gyri it was positive (r(70)=.75, p<.001). Most notably we found an age-related dissociation: whereas the brain-behaviour relation was consistent across age for the angular gyri; Activation in the dmPFC showed a moderation with age F(1,68)=4.19, p=.045). Participants in both adult groups varied in the degree they deactivated the dmPFC, the stronger the deactivation the better the performance. Children showed neither a deactivation of the dmPFC nor a relation to recall performance. Mean memory performance refers to the average across the schema and no-schema paired associates. A star indicates a significance of p<.05.

The two component model helps us to test whether the age-related differences we observed are driven by immaturity of the associative or the strategic component. Memory differences linked to associative regions, such as the angular gyrus, or to the utilisation of schema would indicate differences in the associative component. Memory differences that are not linked to the associative memory regions but rather to regions involved in executive function would suggest a stronger role for the strategic component. To corroborate the links between task activation and executive function we used the independently acquired WCST performance as a general measurement of executive function (Greve et al., 2005): Participants with high levels of executive function in the WCST can likely use those functions strongly to facilitate their retrieval performance.

Children had a lower retrieval performance than adolescents and adults for the schema and the no-schema new paired associates. We found for both the adolescents and the adults, the level of the deactivation of the dmPFC during trials in which they recalled the correct location predicted their overall recall performance: the stronger the deactivation was, the better was the performance (Fig. 3). This deactivation also predicted the WCST performance. Furthermore, the dmPFC cluster we found overlaps with a core cluster previously observed during performance of the WCST (Specht et al., 2009) and is also contained within the Executive Control Network, a resting state network that is involved across many aspects of executive function (Smith et al., 2009). The link to the WCST, the involvement of our dmPFC cluster in the WCST and the dmPFC’s important role in executive function (Ridderinkhof, 2004; Domenech and Koechlin, 2015), suggests to us that it reflects executive function benefitting retrieval performance. Participants with strong deactivation in the dmPFC could use executive function to improve their task performance, whereas participants that showed little or no deactivation could not. In contrast to the other groups, children did not seem to exhibit this behaviour: they neither showed a systematic deactivation of the dmPFC nor was the dmPFC activity related to memory or WCST performance, in which children performed worse than adolescents and adults. We take all this as an indication that the strategic component in children is not as mature as in adolescents and adults: Whereas adolescents and adults can use their strategic abilities to enhance their memory performance, children did not seem to be able to do this. These results are nicely in line with recent work demonstrating that age-related increases in mnemonic strategies is linked to the development of the PFC (Yu et al., 2018).

With regard to the associative component, we did not find any indications for differences between age-groups. Activity of the angular gyrus was correlated with successful memory performance consistently across groups. Additionally, schema effect was indistinguishable across groups. All groups performed between 20 and 30 percent better in the schema over the no-schema condition. We interpret this absence of any developmental differences for associative processes as an indication for a weaker role of the associative component to explain age-related memory differences in our sample.

The consistent relation of the activation from the angular gyrus across groups suggests an important role in the task that is stable across the tested ages. This stability is consistent with previous work demonstrating that the angular gyrus has the same functional boundaries in school children (7 to 10 years old) compared to adults (Barnes et al., 2012); suggesting a relative early functional maturation. In recent years, the contribution of the angular gyrus to memory has received increased attention. There is now substantial evidence for it being an amodal convergence zone (Bonnici et al., 2016; Yazar et al., 2017) that integrates input from different modalities to create higher level representations. With this facility it lays the basis for abstract representations and thus semantic memory (Binder and Desai, 2011). The ability to combine several modalities seems ideally suited for the memory game task where spatial information (location) needs to be combined with semantic information (identity of the card). Another capacity of the angular gyrus that explains its involvement is the ability to guide attention during memory retrieval relying on retrieval cues or recovered memories (Cabeza et al., 2008; Vilberg and Rugg, 2008).

We replicated that schemas facilitate memory (Tse et al., 2007; van Kesteren et al., 2010; van Buuren et al., 2014; Liu et al., 2017) as indicated by the higher performance for the schema new paired associates. This effect did not show any differences across development, in line with a previous study investigating children in a similar age range (Brod et al., 2016). Neurally, we found that not the mPFC but the angular gyrus distinguished the retrieval of schema versus no-schema associations. Both the angular gyrus and the mPFC were activated in both the schema and no-schema condition, however the angular gyrus was significantly more strongly activated whereas the activity of the mPFC did not differ significantly. This pattern is consistent with results previously found using this paradigm (van Buuren et al., 2014), but it appears at odds with the typical pattern that the mPFC orchestrates the utilisation of schemas (van Kesteren et al., 2012; Fernández, 2017; Genzel and Battaglia, 2017). We speculate that the mPFC did not differentially activate as there were too many associations at the same time that needed to be assimilated in the schema. If either there would have been less associations to learn or there would have been more time for learning the associations and stabilising their memories, we speculate that the mPFC would have been more strongly activated for the correctly retrieved schema new paired associates.

In summary, we investigated whether memory differences between children, adolescents and adults would stem from developmental changes in executive abilities, the strategic component, or rather from differences in mechanisms related to binding different features together into a memory representation, the associative component. We found that adolescents and adults outperformed children in memory. The performance within the adolescents and adult group was associated to their individual executive abilities, thus providing evidence that a maturation of the strategic component was driving the age-related differences we observed. In contrast, we did not find differences in the associative component that help to explain the differences in memory between the age groups.

## 5. Acknowledgement

This research was funded by an NWO Research Talent Grant (406-13-008) to N.C.J.M. and G.F.

